# The amyloid packing difference: a pairwise comparison metric for amyloid structures

**DOI:** 10.64898/2026.02.18.706523

**Authors:** Sjors H.W. Scheres

## Abstract

Several proteins from the human proteome have been observed to adopt multiple distinct amyloid filaments, and specific protofilament folds are associated with different diseases. Thereby, it has become necessary to compare pairs of amyloid structures of a given protein. This paper describes the amyloid packing difference (APD), which quantifies the difference between such a pair as the percentage of residues that are involved in unique cross-β packing interactions or that have different side chain orientations relative to the β-strands. Clustering of α-synuclein protofilament folds on pairwise APD values recapitulates previously reported clustering based on structural superpositions. Any pair of known protofilament folds of the prion protein, tau, α-synuclein, TDP-43 or TAF15 from different neurodegenerative diseases have APD values above 20%, whereas all pairs of structures that have been associated with the same disease have APD values below 40%. Structures of antibody light chain from different individuals with systemic amyloidosis differ from each by APDs above 60%, whereas structures of transthyretin filaments from different individuals are strikingly similar, with APDs below 25%. These observations provide context for the interpretation of APD values in future structure comparisons.

## Introduction

The amino acid sequences of proteins have evolved to adopt a specific protein fold that is intrinsically linked to their function. Although some proteins adopt more than one globular fold ^1^, most proteins adopt only a single fold. This principle does not extend to amyloid filaments. Advances in helical reconstruction of electron cryo-microscopy (cryo-EM) images ^2,3^ have enabled the determination of atomic structures for hundreds of amyloid filaments ^4^. Combined, these structures have revealed that a single protein sequence may adopt many different amyloid folds.

Amyloid filaments represent an aggregated state in which multiple copies of the same protein assemble into helical filaments. Each layer of these filaments consists of one or more copies of the protein, which form long twisted β-sheets in the direction parallel to the helical axis. Thereby, each layer is separated from the next by the distance between β-strands in a β-sheet, which is approximately 4.7-4.8 Å. Most known amyloid filaments comprise multiple copies of the same protein, but amyloid filaments that contain two different proteins have been described ^5^. When there are multiple copies of a protein on each layer, the filament is comprised of multiple protofilaments. Either within or in between the protofilaments, β-sheets pack against each other through so-called cross-β packing, which involves interactions between the side chains that stick out from the β-sheets. The ability of a given protein to adopt multiple different amyloid folds arises from flexibility in the nature of the cross-β packing interactions, which can be charged, hydrophilic or hydrophobic.

Among the most studied proteins that form amyloid filaments are those that are implicated in neurodegenerative disease. Only a few proteins from the human proteome have been observed to form amyloid filaments in the brain, and the aggregation of each of them is characteristic of multiple diseases that affect distinct brain regions and/or cell types. Cryo-EM structures of tau, α-synuclein, TDP-43 and TAF15 filaments extracted from *postmortem* brains of individuals with these diseases have revealed that specific amyloid protofilament folds characterise distinct diseases ^6,7^.

In order to understand why specific amyloid folds form in different circumstances, and what their role is in disease, it is imperative to replicate amyloid filaments with disease-specific structures in the laboratory ^6^. Because a given protein may adopt many different amyloid structures, it will be important to quantify the resemblance of one structure to another to measure how successful these efforts are. Methods based on local topology have been explored to identify local regions of interest in amyloid folds ^8^. More often, to compare two structures, one superposes them and reports root mean square deviations (RMSDs) between C-alpha (CA) atom positions or template modelling (TM)-scores ^9^. For example, three groups have recently reported independent classifications of amyloid protofilament folds, either based on CA RMSDs ^10,11^, or on TM-scores ^12^. However, structural superposition may vary for sub-structures, and overall structure matching scores have thus far not prevented the use of ambiguous language to describe the similarity between amyloid folds in the literature.

Here, I propose a new metric for the pair-wise comparison of amyloid structures: the amyloid packing difference (APD). Based on the observation that distinct amyloid folds are driven by specific cross-β side chain packing interactions, the APD directly compares lists of these interactions for any pair of amyloid structures. Thereby, the APD is invariant to the relative orientation of the structures, precluding the need for their superposition. I will show that clustering of α-synuclein protofilament folds based on pairwise APD values recapitulates their classification based on RMSDs or TM-scores and structural superposition. In addition, I will analyse APD values between amyloid structures that are known to be associated with distinct neurodegenerative diseases. This analysis provides context for any comparison of amyloid structures, as I will illustrate for several recently reported structures.

## Approach

The calculation of the APD for any pair of amyloid structures consists of a two-step procedure, each of which is implemented as a separate python script. A third script is auxiliary.

### Step 1: calculate contacts for an amyloid structure

The first script, `contacts.py`, generates a list of contacts for a given input structure. Because the main chain within a given layer of the amyloid filament may meander up and/or down the helical axis (which is assumed to be along the Z-direction), the closest contacts between pairs of residues may be between different layers of the amyloid. The input structure needs to contain sufficient layers to capture all these contacts. To facilitate the generation of structures with multiple identical layers, an auxiliary python script called `helix.py` was implemented to apply helical symmetry operators to a specified subset of chains of an input structure.

The `contacts.py` script identifies protofilaments in the input structure based on the distances between CA atoms of residues with identical residue numbers in the Z-direction. For each protofilament, it then identifies the chain corresponding to the middle layer. For every residue in the middle chain of each protofilament, the script identifies pairwise residue-residue contacts. Two residues are considered to form a contact if at least one pair of their non-hydrogen side-chain atoms (or CA atoms in the case of glycine) are within 6.5 Å. When multiple atom pairs between the same two residues satisfy this criterion, only the shortest distance is stored. Contacts between residues that are adjacent to each other, or separated by one residue within the same chain, are excluded. Moreover, to avoid counting contacts between side chains that are on opposite sides of the protein backbone, the script defines a reference plane for each residue. This plane is defined by the positions of the N and C atoms of that residue, and the position of the N atom of the corresponding residue in the next layer. Then, contacts are only kept if, for both of its residues, the two contacting atoms are on the same side of the plane as its CB atom. For glycines, which lack a CB atom, the N-C-N planes are ignored.

Contacts are classified as intra- or inter-protofilament, depending on whether the two residues involved belong to the same protofilament. To keep track of contacts between different layers of the amyloid, for each contact the script also stores a ca_offset value, which is the difference in Z-coordinates of the CA atoms of the two residues in the middle layer of the amyloid, divided by the average (∼4.75 Å) distance between layers within the amyloid.

Finally, the `contacts.py` script calculates for each residue in the middle layer of the input amyloid structure whether its CB atom is on one side or the other side of its N-C-N plane. The information about all contacts and the orientation of all middle-layer residues with respect to their N-C-N plane is written in a CSV file.

### Step 2: compare lists of contacts for two structures

The second script, `compare.py`, takes the CSV files from two different amyloid structures as inputs to perform the comparison. The two input structures may differ in the extent of the ordered filament core. The script will calculate the number of residues that are ordered in both input structures (*N*_*common*_), as well as the number of residues that are only present in the input structure with the largest ordered core (*N*_extra_).

All contacts in the input CSV files that are between residues that are ordered in both input structures are classified to be either unique to one of the input structures, or in common between the two input structures. Contacts are considered as common contacts if they occur between the same two residues in both input structures, with a distance between non-hydrogen side-chain atoms (or CA atoms for glycines) below 4.5 Å in one of the input structures and below 6.5 Å in the other input structure. The more relaxed distance cutoff for the other input structure provides robustness to side-chain conformations in the calculation of common contacts. Contacts are considered as unique contacts if they have a distance below 4.5 Å in one structure and they are not present in the other structure.

The APD is based on *N*_*different*_, which is the number of residues that are ordered in both structures and that participate in a unique contact and/or have a different orientation with respect to their N-C-N plane between the two input structures. The APD is then calculated as:

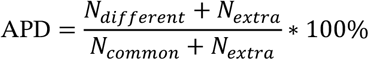

The *N*_*extra*_ terms in the calculation of the APD will penalise structures with distinct extents of their ordered cores, effectively considering all *N*_*extra*_ residues as different. The `compare.py` script outputs two different distances: for the XY-APD it calculates the number of residues with unique contacts (*N*_*uniqueXY*_) by ignoring whether contacts happen between the same or different amyloid layers. For the XYZ-APD, the script also considers whether residues have the same offsets in the Z-direction to calculate the number of unique contacts (*N*_*uniqueXYZ*_). The Z-offsets are integer values calculated from the Z coordinates of the CA atoms, rounded to multiples of 4.75 Å. Because the XY-APD ignores the Z-offsets, the XYZ-APD of a given comparison cannot be lower than its XY-APD. APD values can be calculated for the comparison of individual protofilament folds, as well as for inter-protofilament interfaces. In the latter case, only residues involved in contacts between protofilaments in both structures are considered.

The `compare.py` script calculates XY-APD and XYZ-APD values for all protofilaments and inter-protofilament interfaces in the input structures. In addition, it generates a schematic representation of the comparison. Residues that are in common among the two input structures are represented by a white circle with a black outline and its one-letter amino acid code in black; residues that are only present in one of the structures are represented with a white circle with a grey outline and a grey one-letter code; connections between the circles represent the main chain. The circles of residues with different side chain orientations relative to their N-C-N plane between the input structures are filled orange; the circles of residues that are mutated between the two structures are filled red. Contacts that are in common between the two input structures are shown in light grey lines; contacts that are unique are shown in dark orange for intra-protofilament contacts and in light orange for inter-protofilament contacts; contacts that are only unique due to differences in Z-offsets are shown in marine; and contacts that involve residues that are only present in one of the input structures are shown in dark yellow.

## Results

### A few illustrative examples

Among the proteins that form amyloid filaments in disease, the prion protein is probably the best characterised in terms of its effects on disease. Prion strains that cause distinct forms of prion disease have been isolated and propagated in the laboratory for several model organisms ^13,14^. **Figure 1A** shows the structures of two mouse-adapted strains: PDB entry 7QIG is the mouse aRML strain ^15^; PDB entry 8EFU is the mouse a22L strain ^16^. The structures were superposed using matchmaker in UCSF Chimera ^17^ on residues 94-225, which are in common between their ordered cores. The RMSD between their superposed CA atoms is 4.30 Å. Visual inspection suggests the presence of a hinge movement between the lower N-terminal lobe and the upper C-terminal lobe. Superposition on residues 94-165 gives an RMSD of 2.37 Å; superposition on residues 166-225 gives an RMSD of 4.84 Å (**Figure 1B-D**). These values suggest that the lower N-terminal lobes are similar to each other, while the upper C-terminal lobe is less conserved between the two structures.

**Figure 1:**
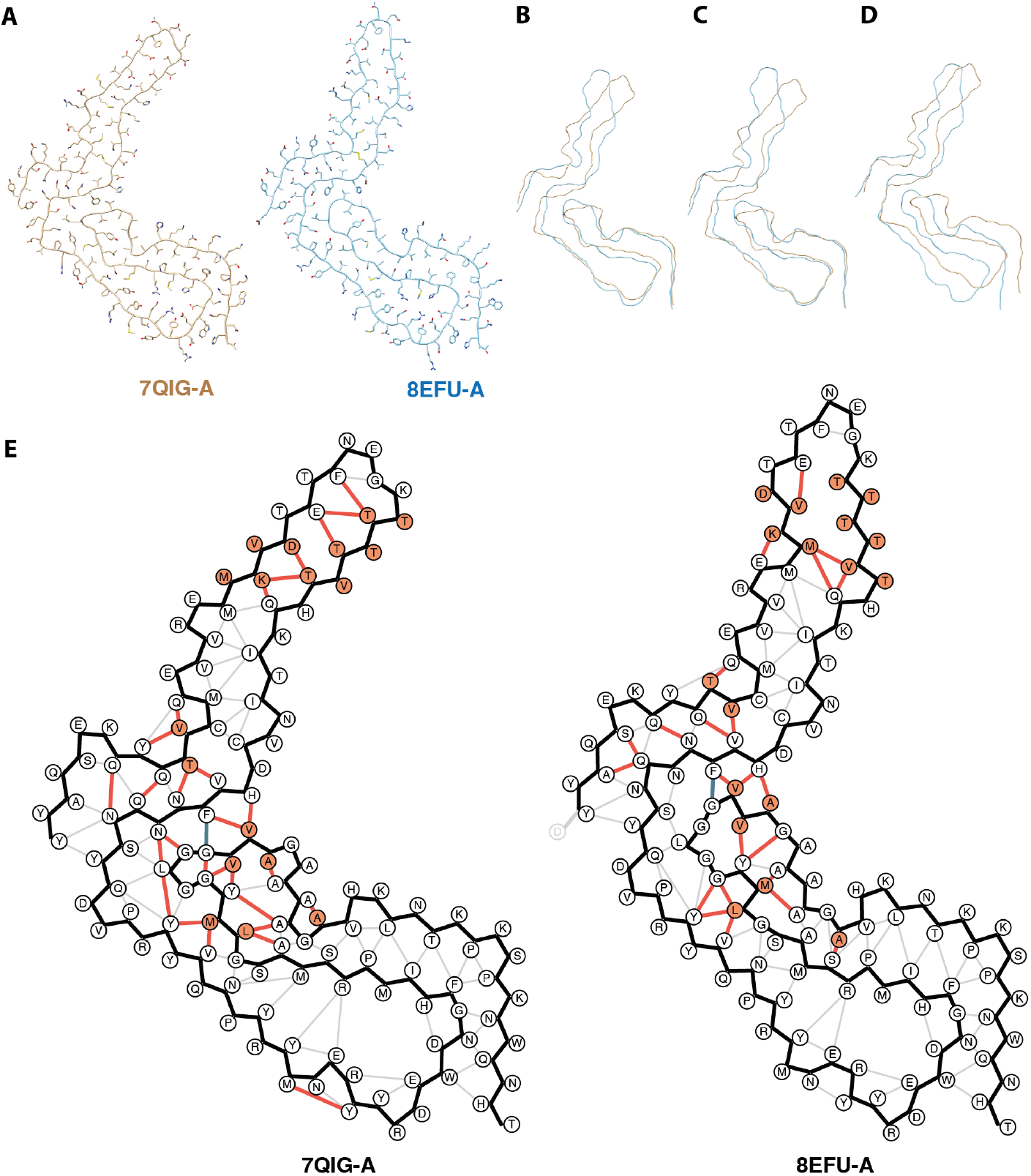
An illustrative example. **(A)** Stick models of the atomic structures of chain A from PDB entry 7QIG (brown) and of chain A of PDB entry 8EFU (light blue). Backbone CA-trace of the two structures from panel A, superposed on residues 94-225 **(B)**, 94-165 **(C)** or 166-225 **(D)**. (**E)** Schematic representations of the structures in A as output by the compare.py script. The main chain is shown with black lines; residues with a side chain flip relative to their N-C-N planes in orange circles; contacts between residues with dark orange lines; common contacts without a Z-offset with grey lines; and common contacts with a Z-offset in teal green lines. For this comparison: *N*_*common*_=132, *N*_*extra*_=1, *N*_*flips*_=18, *N*_*uniqueXY*_=*N*_*uniqueXYZ*_=48, and XY-APD=XYZ-APD=38%.

These observations illustrate one of the problems with pairwise structure superposition and the use of RMSD for structure comparison. If overall rotations exist between separate parts of a structure, then the overall RMSD may not reflect resemblance between the individual parts. However, the division of a structure into separate parts for pairwise superpositions is often not straightforward, especially in automated procedures. The APD catches the similarity of amyloid sub-structures without the need for their separate superposition.

**Figure 1E** was generated from the two PDB files using the following commands:

~~~
python helix.py 7QIG.pdb --chains A --twist -0.64
--rise 4.82 --box_and_pixel 384 1.067 --n_copies 4
--output 7QIG-A.pdb
python helix.py 8EFU.pdb --chains A --twist -0.565
--rise 4.75 --box_and_pixel 384 1.045 --n_copies 4
--output 8EFU-A.pdb
python contacts.py 7QIG-A.pdb
python contacts.py 7LNA-A.pdb
python compare.py 7QIG-A_contacts.csv 8EFU-A_contacts.csv --flip1 --flip2 --rot1 180 --rot2 70
~~~

The `helix.py` script applies helical symmetry to chain A of each PDB file using the supplied twist (in degrees) and rise (in Å). The --n_copies argument generates 4 additional copies of the chain above and below the original chain. If the atomic model was built into a map generated by helical reconstruction in RELION ^3^, the --box_and_pixel argument calculates the XY coordinates of the helical axis from the box size of the map (in pixels) and the pixel size (in Å). Alternatively, the helical axis coordinates may be specified using the --axis_xy argument. The `contacts.py` script requires no arguments, other than the coordinate file generated by the `helix.py` script. The `compare.py` script includes arguments to control the orientation of the schematic representation (--flip1 and --flip2), while the --rot1 180 and --rot 70 arguments apply clockwise rotations of 180° and 70° to the respective representations. Note that the mirror and rotation operations of the script do not affect the APD calculation (which is rotation and mirror invariant) and are merely used to orient the schematic representations with the structures shown in Figure 1A-D for convenience of visual comparison.

**Figure 1E** gives insights into the similarities and differences between the two structures. For example, flips in the side chains of methionine 128 and leucine 129 relative to their N-C-N planes in the middle layer of the lower N-terminal lobe leads to unique packing contacts in both structures. Unique contacts that arise from similar flips of alanine 112 and of alanine 119, valine 120 and valine 121 also give rise to changes in the N-terminal lobe. In addition, unique interactions between the latter triplet change the packing of the lower lobe to the upper lobe. Side chain flips relative to their N-C-N planes for multiple residues at the tip of the upper lobe give rise to distinct unique contacts there, whereas flips of valine 214 and threonine 215 give rise to unique contacts in its lower base. The packing of the last 10 residues of the ordered core against the lower domain involved unique contacts between the two structures that are not caused by such flips.

Both the XY and the XYZ variants of the APD are 38%. This is because there is only a single contact with the distinct Z-offset between the two structures, between glycine 122 and phenylalanine 174, and both residues are also involved in other unique contacts. Of the 132 residues that are in common between the two structures there are 18 residues with side chain flips relative to their N-C-N plane (*N*_*flips*_) and 48 residues that are involved in unique contacts; there is only one extra residue (aspartate 226 in the a22L strain).

**Supplementary Figure 1** shows the schematic representation of the compare.py script for the comparison between the aRML mouse strain mentioned above, and a second structure of the same mouse strain, 7TD6, that was determined independently by another group. 7QIG was determined to a resolution of 2.7 Å ^15^; 7TD6 was determined to a resolution of 3.0 Å ^18^. Despite the reasonably high reported resolutions, the two structures do show some differences. The XY-APD is 8%; the XYZ-APD is 16%. The XY differences arise from contacts that are identified as unique between methionine 143 and tyrosine 146 in 7QIG; and between alanine 168 and tyrosine 223 in 7TD6. These differences are caused by different side chain orientations in the two structures, leading to distance pairs beyond 6.5 Å. This sensitivity of the APD to side chain rotamers is difficult to avoid and illustrates how some contacts that are being identified as unique may not contribute to different amyloid folds in practice. In the analyses performed below, such differences typically lead to noise in the APD values of a few percent.

A more systematic difference is present in the tip of the C-terminal lobe, where 9 contacts between 11 residues have a different Z-offset. Inspection of the atomic models shows that these Z-offsets arise from different tracings of the main chain: from isoleucine 181 until threonine 191 on one side of the arm, the main chain paths of the two structures deviate, which results in the measured Z-offsets and the increased XYZ-distance. Inspection of the cryo-EM maps reveals that the map of the 7TD6 structure shows spurious connections between the different amyloid layers (**Supplementary Figure 2**). At a resolution of 3.0 Å, such connections should not be present. They may thus be an indication of the reconstruction getting stuck in a local minimum, where projections of an incorrect reconstruction still represent a reasonable fit to the experimental data ^19^. Therefore, it remains unclear whether this difference represents a real difference or whether it arose from a suboptimal reconstruction. Because reconstructions with spurious connections between the amyloid layers are relatively common, and they may result in artifacts in the XYZ-APDs, the analyses in the remainder of this paper were all performed using the XY-APD. As microscopes, electron detectors and image processing programs continue to improve, the problems with amyloid reconstructions getting stuck in local minima will become less, and the XYZ-APD will suffer less from these artifacts.

To reflect the situation when the residue numbering of the input structures differs (for example because they come from different organisms), the user may specify a residue number offset for the second structure through the --offset2 parameter of the compare.py script. For example, **Supplementary Figure 3** shows the comparison between PDB entries 7QIG and 7LNA, with the latter representing the 263K strain in hamster ^20^. Because threonine 94 in mouse is equivalent to threonine 95 in hamster, the compare.py script was run with the additional argument --offset2 -1. The two structures have an (XY-) APD of 61%, 21 residues out of the 129 residues that are in common among their ordered cores have side chain flips relative to their N-C-N planes; 74 residues are involved in unique contacts; 8 residues are mutated between the mouse and hamster sequences; and the number of extra residues is 3 (arising from three modelled residues at the tip of the long arm in the mouse structure that were unmodelled in the hamster structure).

### The APD recapitulates classifications of α-synuclein proto-filament folds

α-Synuclein is the protein with the highest number of amyloid structures in the PDB. Recently, three groups have reported independent classifications of α-synuclein protofilament folds ^10–12^. Although they use different methods of clustering, all three groups use pair-wise structure superpositions and compare pairs of protofilaments based on CA RMSDs ^10,11^ or based on TM-scores ^12^. Milchberg et al identify two main clusters and 14 folds that are distinct from the structures in these clusters and remain unclassified. They then subdivide the first main cluster into two sub-clusters (called 1A and 1B) and the second main cluster into three subclusters (2A, 2B and 2C). Connor et al classify α-synuclein folds into 11 groups, with most folds pertaining to groups 4 or 8. The classification by Price et al is based on a TM-score cutoff of 0.5, which results in 14 groups, with the two largest groups, as well as the smaller groups, closely matching those of Connor et al.

**Figure 2A** shows a hierarchical clustering (with median linkage) of α-synuclein protofilament folds based on all pairwise XY-APD values. Cluster 2 from Milchberg et al (yellow, orange and purple squares) overlaps with group 4 from Connor et al (purple dots) and cluster 2 from Price et al (purple triangles). Cluster 2 separates from the rest of the structures at an APD above 70%. Subclusters 2A (yellow squares), 2B (orange squares) and 2C (purple squares) from Milchberg et al also separate in the APD-based clustering, albeit at lower values (23-32%). Cluster 1 from Milchberg et al (blue and green squares) overlaps with groups 7 (orange dots) and 8 (red dots) from Connor et al, where structures from group 7 are all part of subcluster 1B (blue squares), and with cluster 1 (red triangles) from Price et al. APD-based clustering separates clusters 1A and 1B at an APD just above 50%. APD-based clustering also identifies structures 8H05 and 7V49 as distinct from the other structures in cluster 1, which agrees with the separation of these structures in Figure 4A by Milchberg et al. All structures that are unclassified by Milchberg et al, as well as all groups with at most 3 structures by Connor et al, or in groups with at most 5 structures by Price et al, separate at ADP values above 50%. The excellent agreement between the APD-based clustering and the three independent classifications that are based on pair-wise structure superpositions provides support for the underlying assumption of the APD: that distinct amyloid folds arise from different contacts between side chains of residues in cross-β packings.

**Figure 2:**
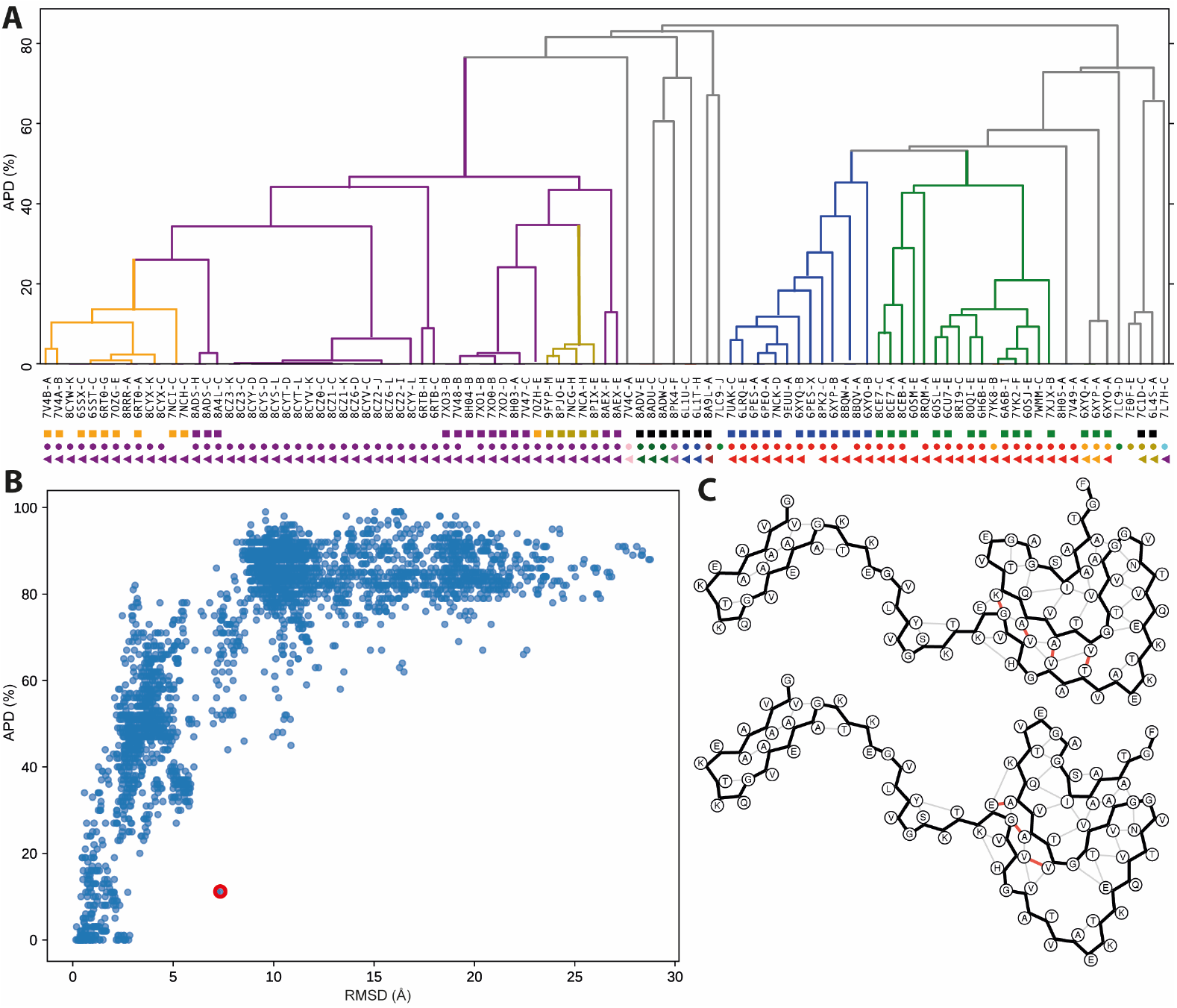
Clustering of α-synuclein protofilament folds based on the APD. **(A)** PDB codes and chain IDs are shown for all structures that are clustered. Coloured squares represent the clustering by Milchberg et al; dots represent the clustering by Connor et al; and triangles represent the clustering by Price et al. The squares are coloured like the clusters by Milchberg et al, and parts of the dendrogram are coloured with the same colours for convenience of comparison. The dots are coloured like the clusters in the paper by Connor et al. Triangles are coloured in the same colours as the dots, to reflect the similarity in clustering by Connor et al and Price et al. **(B)** Scatter plot of APDs versus RMSDs of the structures clustered in panel A and present in the data set by Connor et al. **(C)** Example of a pair with a relatively low APD (11%) and high RMSD (7.18 Å): chains A of 6XYO and 6XYP, as indicated with a red circle in panel B.

To further analyse how APD relates to the RMSD of aligned α-synuclein protofilament folds, **Figure 2B** shows a scatter plot of the APD versus the RMSD as calculated by Connor et al. This plot confirms the correlation between the APD and the RMSD, where APDs tend to rise from 0 to 75% when RMSDs increase from 0 to 5 Å. Almost all structures with APDs higher than 80% have RMSDs larger than 8 Å. **Figure 2C** shows an exception to these overall trends (indicated with a red circle in Figure 2B): for protofilaments A of MSA type I and type II filaments, respectively. These proto-filaments have an APD of only 11%, despite having an RMSD of 7.18 Å. The relatively low APD reflects the observation that two near-identical substructures (glycine 1 to lysine 30 align with an RMSD of 0.26 Å; glutamate 48 to phenylalanine 81 align with an RMSD of 0.61 Å) pack against each other through interfaces that involve relatively few residues.

### APDs for structures with known pathological outcomes

Specific protofilament folds of proteins that form intracellular lesions have been observed to characterise distinct diseases ^6^. **Figure 3** shows hierarchical clustering of protofilament folds of the prion protein, tau, α-synuclein, TDP-43, antibody light chain and transthyretin based on (XY-) APD values between corresponding brain-derived amyloid structures in the Amyloid Atlas. The clustering of recently reported TAF15 structures ^7^ is also shown. Across all five proteins, clustering based on APD values recapitulates classifications based on the associated diseases.

**Figure 3:**
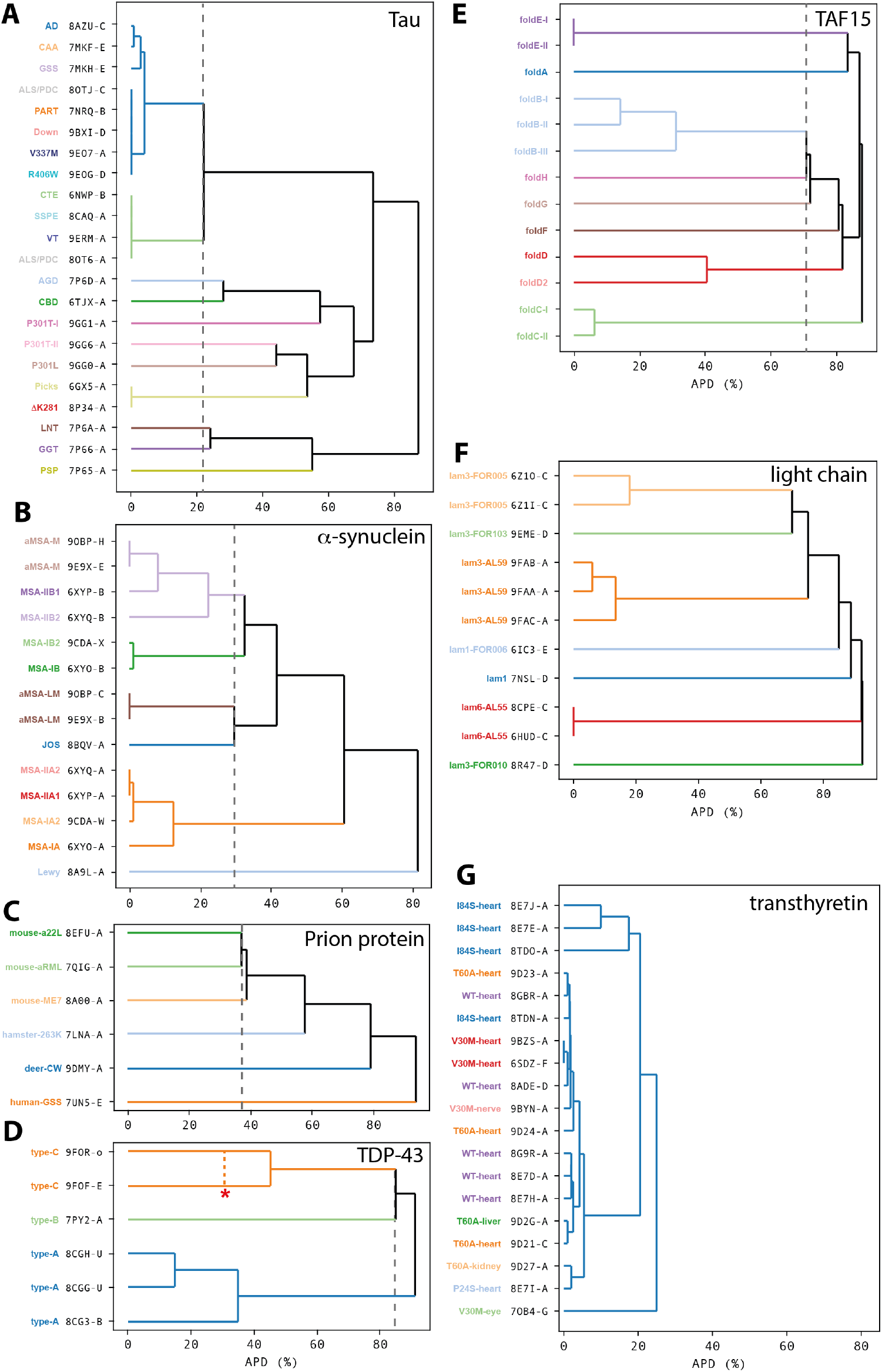
Clustering of α-synuclein protofilament folds based on the APD. **(A)** PDB codes and chain IDs are shown for all structures that are clustered. Coloured squares represent the clustering by Milchberg et al; dots represent the clustering by Connor et al; and triangles represent the clustering by Price et al. The squares are coloured like the clusters by Milchberg et al, and parts of the dendrogram are coloured with the same colours for convenience of comparison. The dots are coloured like the clusters in the paper by Connor et al. Triangles are coloured in the same colours as the dots, to reflect the similarity in clustering by Connor et al and Price et al. **(B)** Scatter plot of APDs versus RMSDs of the structures clustered in panel A and present in the data set by Connor et al. **(C)** Example of a pair with a relatively low APD (11%) and high RMSD (7.18 Å): chains A of 6XYO and 6XYP, as indicated with a red circle in panel B.

For tau, various structures of paired helical filaments (PHFs) with the Alzheimer fold cluster together, as do structures with the chronic traumatic encephalopathy (CTE) fold. Structures of tau protofilament folds from other tauopathies are clustered in excellent agreement with the previously postulated structure-based classification ^21^. Structures that are associated with different tauopathies separate from each with APD of ∼20% and higher.

Structures of α-synuclein filaments are available for three synucleinopathies. Filaments from multiple system atrophy (MSA) consist of two types and each type has two different protofilament folds A and B. Filaments from juvenile-onset synucleinopathy (JOS) and from individuals with Lewy pathology comprise only a single protofilament. The JOS fold separates from protofilaments B of the MSA fold at an APD of ∼30%; the two protofilament types of MSA separate at an APD of ∼60% and the Lewy fold separates from all other synuclein folds at an APD above 80%.

For the prion protein (also see Figure 1), structures of different mouse strains are more similar to each other (with APDs just under 40%) than to structures of strains from hamster, deer or human (with APDs above 50%).

Structures of TDP-43 filaments from three different TDP-43 proteinopathies are available: types A, B and C. Two of the folds were observed to have two or more variants, which were defined as distinct conformations that co-exist as continuous stretches within individual filaments ^22,23^. The three different types separate at APDs above 80%, whereas the three variants of the type-A folds separate at APD values in the range of 15-35%, while two variants of the type C fold have an APD of 43%. The type C fold is unique among the known TDP-43 filaments, as it contains a second protein, annexin A11, in addition to TDP-43. The packing of annexin A11 is identical between the two type C variants (**Supplementary Figure 4**). When also considering the 36 residues of annexin A11 as part of the common residues, the ADP between the two type C folds is reduced to 31%.

For TAF15, structures are available from eight distinct TAF15 proteinopathies: types A-H. The eight types separate at APDs above 70% and are clustered in agreement with the previously proposed structure-based classification ^7^. Again, variants that co-exist within individual filaments were observed for some of the folds (types B, C and E). These variants separate at APD values of 30% or lower. TAF15 filaments extracted from the brain of one individual with neuronal intermediate filament inclusion body disease (NIFID) adopted primarily fold-D, but 1% of the filaments had an alternative conformation, fold-D’, which differs from fold-D with an APD of 41%. These two folds were not observed to co-exist within individual filaments.

The presence in the brain of amyloid filaments of the proteins in Figure 3A-E are linked to neurodegenerative disease. Panels F and G show APD-based classifications of the protofilament folds of antibody light chain and transthyretin, respectively. Filaments of these proteins are linked to peripheral amyloidoses, most often in the heart. The amino acid sequence of the light chains that form amyloid filaments among different individuals with antibody light chain amyloidosis is variable. Consequently, the protofilament folds of light chain filaments from different individuals differ from each other with APDs above 60%, with fold variants that were classified from filaments of a given individual varying up to 20%. This picture is radically different for transthyretin, where amyloid filaments from different individuals, from different tissues, and from wildtype or various mutants all adopt a strikingly similar fold, with APDs below 25%. The largest differences are caused by distinct conformations of a stretch of eleven residues, from glycine 57 and glycine 67, that has been termed the gate ^24^.

Figure 3 recapitulates known classifications of human diseases and suggests a range of APD values, above which the differences between two amyloid protofilaments folds may be physiologically relevant. For tau, α-synuclein and the prion protein, APD values above 20-30% correspond to different neurodegenerative diseases. For TDP-43 and TAF15, APD values in this range may still correspond to variants of folds. So far, considering all known structures of filaments of these five proteins from human brains, APD values above 41% (between the two TAF15 folds of type D) have always been observed to correspond to different diseases. Whereas filaments from different individuals with antibody light chain amyloidosis differ from each other with APDs above 60%, filaments from different cases of transthyretin amyloidosis are remarkably similar to each other. APD values below 25% suggest that distinct conformations of its eleven gate residues, which have been described as polymorphism ^24^, are probably better described as variants of the same fold.

### Measuring the success of replicating disease-specific amyloid structures

The lack of a quantitative comparison metric for amyloid folds has led to ambiguity in reports of similarity between folds in the literature. **Figure 4** shows three examples how the APD may provide more quantitative insights instead.

**Figure 4:**
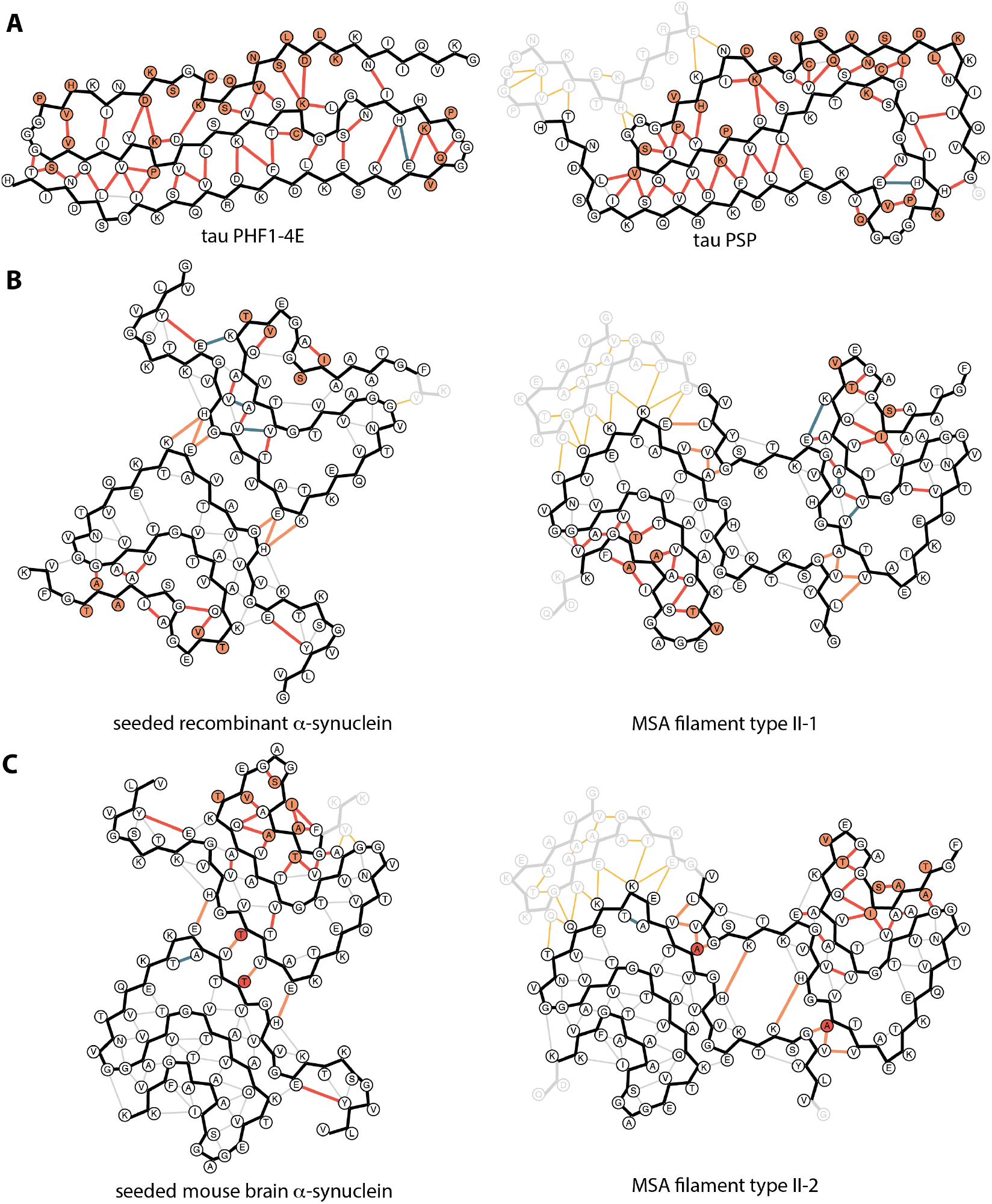
Measuring the success of attempts to replicate disease-specific structures. **(A)** Comparison between spontaneously assembled recombinant tau with four phosphomimetic mutations (tau PHF1-4E; PDB-id 8TTN) versus the progressive supranuclear palsy fold (7P65). For this comparison: *N*_*common*_=90, *N*_*extra*_=20, *N*_*flips*_=27, *N*_*uniqueXY*_=*N*_*uniqueXYZ*_=66, and XY-APD=XYZ-APD=80%. **(B)** Comparison between recombinant α-synuclein assembled by seeded aggregation in vitro (8UKA) and MSA filament type II-1 (6XYP). For the lower left protofilament: *N*_*common*_=61, *N*_*extra*_=3, *N*_*flips*_=5, *N*_*uniqueXY*_=*N*_*uniqueXYZ*_=21, and XY-APD=XYZ-APD=38%. For the upper right protofilament: *N*_*common*_=59, *N*_*extra*_=22, *N*_*flips*_=4, *N*_*uniqueXY*_=23, *N*_*uniqueXYZ*_=24, and XY-APD=56%, XYZ-APD=57%. For the protofilament interface: *N*_*common*_=2, *N*_*extra*_=19, *N*_*uniqueXY*_=*N*_*uniqueXYZ*_=2, and XY-APD=XYZ-APD=100%. **(C)** Comparison between α-synuclein filaments from seeded mouse brains (9RZF) and MSA filament type II-2 (6XYQ). For the lower left protofilament: *N*_*common*_=61, *N*_*extra*_=3, *N*_*flips*_=2, *N*_*uniqueXY*_=2, *N*_*uniqueXYZ*_=4, and XY-APD=8%, XYZ-APD=11%. For the upper right protofilament: *N*_*common*_=58, *N*_*extra*_=23, *N*_*flips*_=7, *N*_*uniqueXY*_=*N*_*uniqueXYZ*_=26, and XY-APD=XYZ-APD=60%. For the protofilament interface: *N*_*common*_=7, *N*_*extra*_=17, *N*_*uniqueXY*_=*N*_*uniqueXYZ*_=7, and XY-APD=XYZ-APD=100%.

The spontaneous assembly of recombinant protein into filaments with the same structures as those observed in human disease is an important goal. This has so far only been achieved for tau, where assembly with truncated constructs or the introduction of twelve phosphomimetic mutations (serine or threonine to aspartate or glutamate) in full-length tau have yielded filaments with ordered cores that are identical to those observed in Alzheimer’s disease ^25,26^. The introduction of four phosphomimetic mutations, around the epitope of the PHF1 antibody, to full-length tau led to the assembly of a three-layered fold that was called PHF1-4E ^27^. This structure was reported to “resemble” the three-layered fold of progressive supranuclear palsy (PSP) ^28^.

However, analysis of the schematic representation in **Figure 4A** shows that the PHF1-4E structure and the PSP structure ^21^ do not have a single cross-β packing interaction in common. The only similarity between the two folds is that the same residues face outwards from the ordered cores from glycine 1 to lysine 8 and from valine 67 to histidine 90. Consequently, the reported APD is 80%. This value is among the highest values at which structures that are specific for different diseases separate in Figure 3, which represents a challenge to the authors’ claim of resemblance.

An alternative method to obtain recombinant filaments with disease-specific folds is by seed amplification, where small amounts of amyloid filaments (the seeds) amplify the assembly of monomeric proteins into new filaments. Although seed amplification is often thought to replicate the structure of the seeds, cryo-EM structures of recombinant α-synuclein with seeds extracted from the brains of individuals with multiple system atrophy (MSA) showed that this is not necessarily the case ^29^. Performing a similar seed amplification assay (SAA), ^30^ reported that “Seed amplification of MSA α-synuclein aggregates preserves the […] structural properties of brain-derived aggregates”. The recombinant filaments and MSA filaments both comprise two protofilaments, but the protofilaments pack symmetrically in the recombinant filaments, whereas two distinct protofilaments pack against each other asymmetrically in MSA filaments.

In the sections above, the `compare.py` script was used to compare individual protofilament folds. **Figure 4B** shows an example of using the script to compare structures with more than one protofilament: for the comparison of the recombinant seeded structure and the type II-1 MSA structure ^31^. The APDs between the recombinant protofilament structure and the structures of protofilaments A and B from the MSA filament are 38% and 56%, respectively. These values are higher than the APD between the JOS fold and the protofilament folds from an atypical case of MSA ^32^. Moreover, the inter-protofilament packing in the asymmetric MSA filaments is completely different from the symmetrical interface in the recombinant filaments, resulting in an inter-protofilament APD of 100%. Therefore, also in this case, the APD values represent a challenge to the authors’ statement that the structural properties of the brain-derived aggregates were preserved in the seed amplification experiment.

Seed amplification can also be performed *in vivo*.

Injection of recombinant α-synuclein filaments into the brains of wildtype mice induces seeded aggregation of mouse α-synuclein, yielding a structure that was reported to “mimic the fold of α-synuclein observed in one protofilament of fibrils isolated from patients with MSA” ^33^. Similar to the seeded recombinant α-synuclein filaments, the filaments formed in the mouse brain consist of two symmetrically arranged protofilaments.

**Figure 4C** shows the output of the compare.py script. The protofilament fold from seeded aggregation in the mouse brain has an APD of 8% with protofilaments A from type II-2 MSA filaments ^31^. Thereby, in this case, the claim that the recombinant fold mimics one of the MSA folds is indeed reflected in an APD value that is lower than APD values that have been observed for variants of folds that co-exist within individual filaments extracted from human brains. The implications of the 60% APD with protofilament B from the same MSA filament, and the 100% APD value for the interprotofilament packing, remain unclear.

## Discussion

This paper introduces a new metric for the comparison of amyloid structures of a given protein: the amyloid packing difference, or APD. Commonly used metrics for structural comparisons, such as RMSDs between atom positions or TM-scores, rely on structural superposition and are therefore difficult to use in cases where parts of a structure adopt distinct relative orientations (e.g. Figure 1B-D). Moreover, RMSDs are often calculated using CA atoms, which can lead to relatively low values for structures that share similar backbone conformations but differ in their side chain packing interactions (e.g. Figure 4B). Because the APD is based on inter-residue side chain interactions and on the orientations of side chains with respect to the main chain, it measures directly the cross-β packing interactions that drive the formation of distinct amyloid folds. In addition, by considering only lists of side chain contacts and orientations, the APD is invariant to the relative orientation of the structures to be compares, which obviates the need for structure superposition. The observation that hierarchical clustering of α-synuclein protofilament folds based on all pairwise APD values recapitulates existing classifications based on RMSD or TM-score between superposed structures validates the APD as a measure for amyloid fold similarity.

Because the APD is based on side chain interactions, it is more sensitive to incorrectly modelled side chains than the backbone RMSD. To increase robustness of the APD to individual side chain orientations, the interatomic cutoff distance for the second member of a contact pair was raised from 4.5 Å to 6.5 Å. Still, as discussed for the example of two independently determined structures of the aRML mouse prion in Supplementary Figure 1, some common contacts may still be identified as unique. Increasing the second cutoff distance risks becoming insensitive to differences that are real, for example the change in the orientation of tyrosine 39 between protofilaments with the Lewy fold, when comparing wildtype and mutant A53T α-synuclein ^34^. In general, and in line with previous recommendations ^19,35,36^, to reduce the risk of introducing errors, atomic models built in cryo-EM maps of amyloid filaments with resolutions worse than 4 Å, or in maps that do not show the expected density features for the given resolution, should be considered unreliable and therefore not be used for APD calculation.

Although specific amyloid protofilament folds of a given protein are associated with different diseases, the pathological relevance of differences among amyloid structures remains unknown. It could be that the different structures merely correlate with the different diseases, or it could be that different structures have specific effects on the cell. Different (stereo)chemistries of the filament surfaces could lead to distinct specific interactions with cellular components, such as cellular uptake receptors and molecular chaperones, or with its own fuzzy coat. It may also take varying amounts of energy to turn over filaments with different folds, owing to distinct stabilities ^37^. Moreover, filaments with different inter-protofilament packings have been observed within the brains of individuals with a given disease, and their different packing interfaces may also lead to distinct interactions with cellular components or different stabilities in the cell. Until the molecular mechanisms by which amyloid filaments affect the cell become clear, the assignment of pathological relevance based on structural information alone will remain difficult.

Still, it may be helpful to interpret APD values for new comparisons in the context of APD values for structures that are known to be associated with different diseases (Figure 3), for example to analyse the extent to which experimental variables, like pH, ionic strength, mutations, and post-translational modifications, influence the reconstitution of disease-specific filaments in model systems of filament formation. For all known structures of brain-derived amyloid filaments of the prion protein, tau, α-synuclein, TDP-43 and TAF15, structures from different diseases have APD values above 20%. Likewise, all structures that have been associated with a given disease have APD below 40%. Therefore, I would argue that two structures should not be claimed to be similar if they have APDs above 40% (e.g. Figure 4A); such claims are probably justified at APDs below 20% (e.g. Figure 4C); and values in between require nuance. Moreover, the schematic representation that is output by the compare.py script provides a convenient way to visualise the differences between two amyloid structures, and this image should be included in reports that use the APD.

## Acknowledgements

I am grateful to Michel Goedert, Sofia Lövestam, Alexey Murzin and Benjamin Ryskeldi-Falcon for critical comments on the manuscript, and to Jake Grimmett, Toby Darling and Ivan Clayson for support with computing. This work was supported by the Medical Research Council, as part of the U.K. Research and Innovation (UKRI) (MC_UP_A025_1013 to S.H.W.S.).

## Code availability

The code is distributed under the (open-source) MIT license and is freely available from http:/github.com/3dem/APD.

## Copyright statement

For the purpose of open access, the MRC Laboratory of Molecular Biology has applied a CC BY public copyright licence to any Author Accepted Manuscript version arising.

---

**Supplementary Figure 1:**
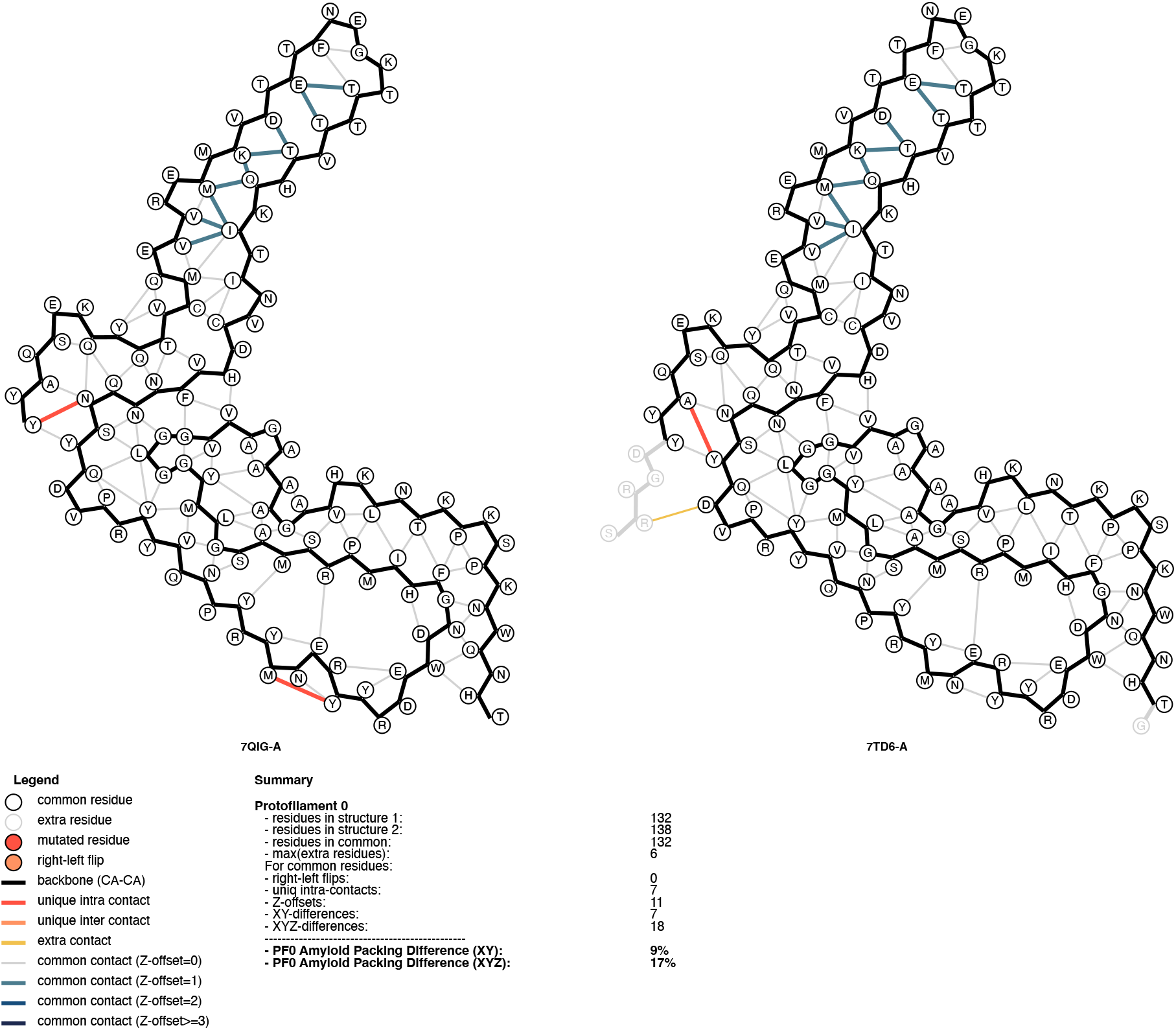
Output of the compare.py script for two independently determined structures of the mouse prion strain aRML (PDB entries 7QIG and 7TD6).

**Supplementary Figure 2:**
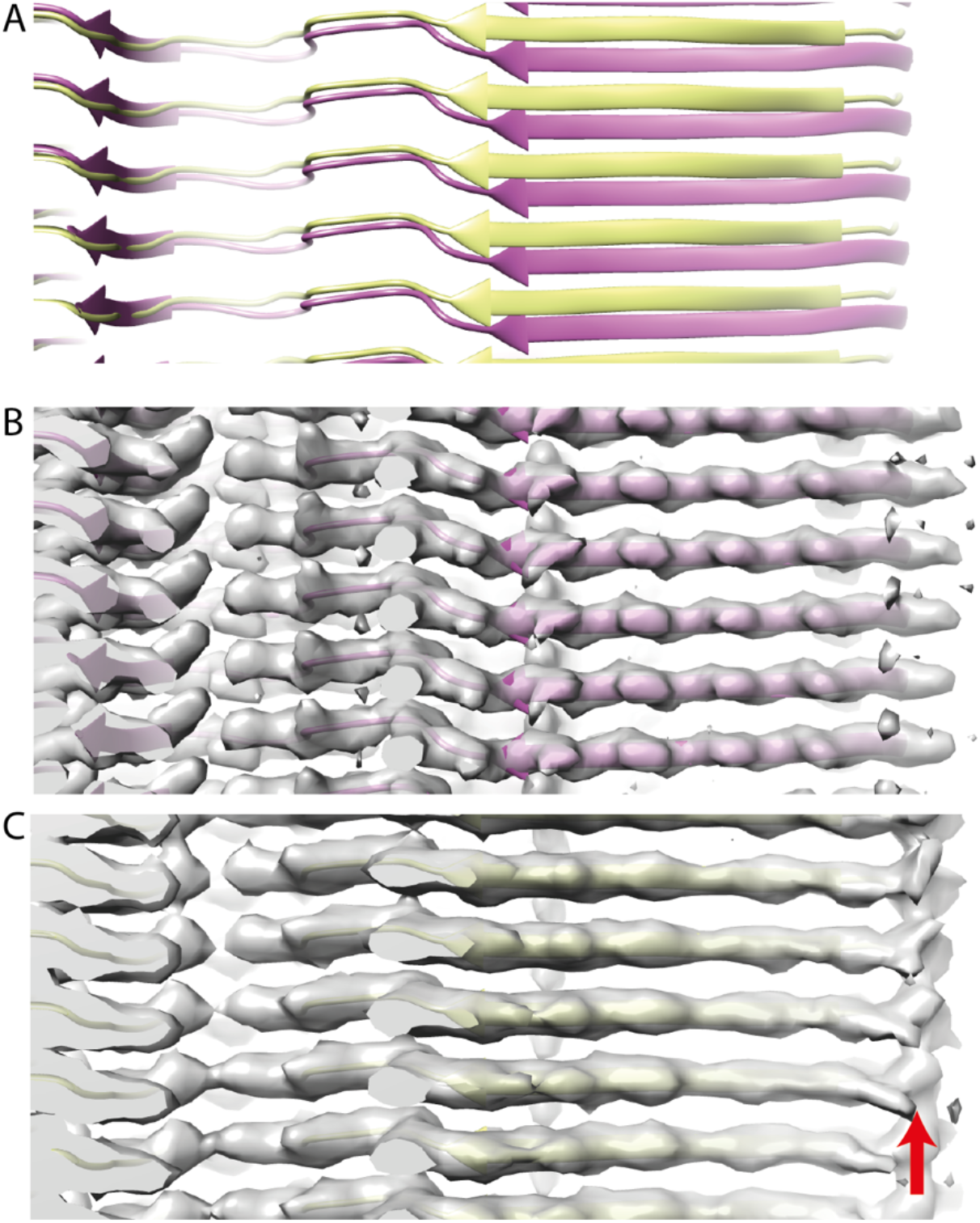
Different Z-tracing in the C-terminal lobe of two independently determined cryo-EM structures of mouse prion strain aRML. **(A)** Superposed main-chain trace of PDB entries 7QIG (pink) and 7TD6 (yellow). **(B)** Main-chain trace and cryo-EM density of structure 7QIG. **(C)** As B, but for 7TD6. The red arrow indicates spurious connecting densities between layers of the amyloid that are indicative of the reconstruction being the result of a refinement getting stuck in a local minimum.

**Supplementary Figure 3.**
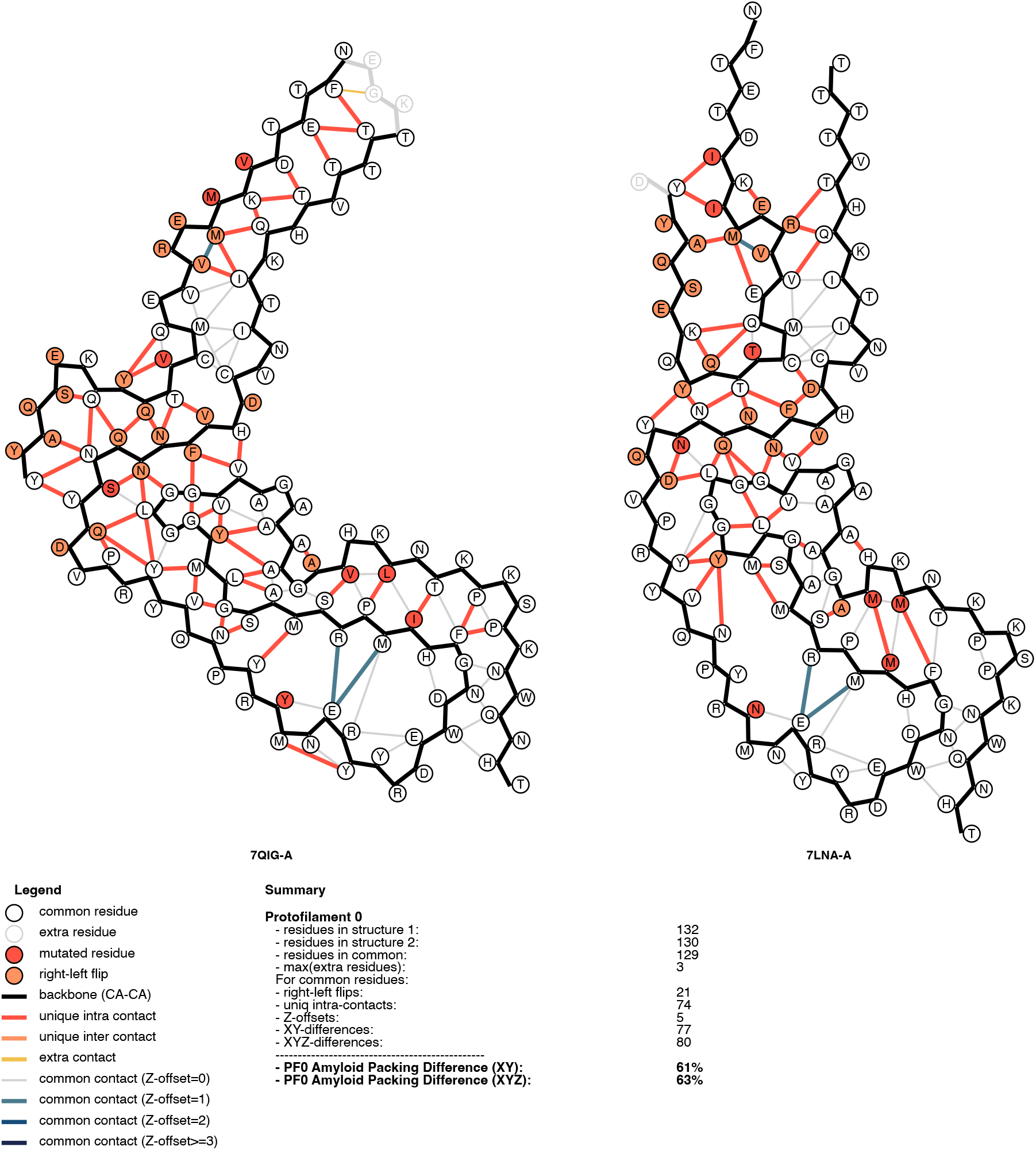
Output of the compare.py script for the mouse prion aRML strain (PDB entry 7QIG) and the hamster prion 263K strain (PDB entry 7LNA).

**Supplementary Figure 4:**
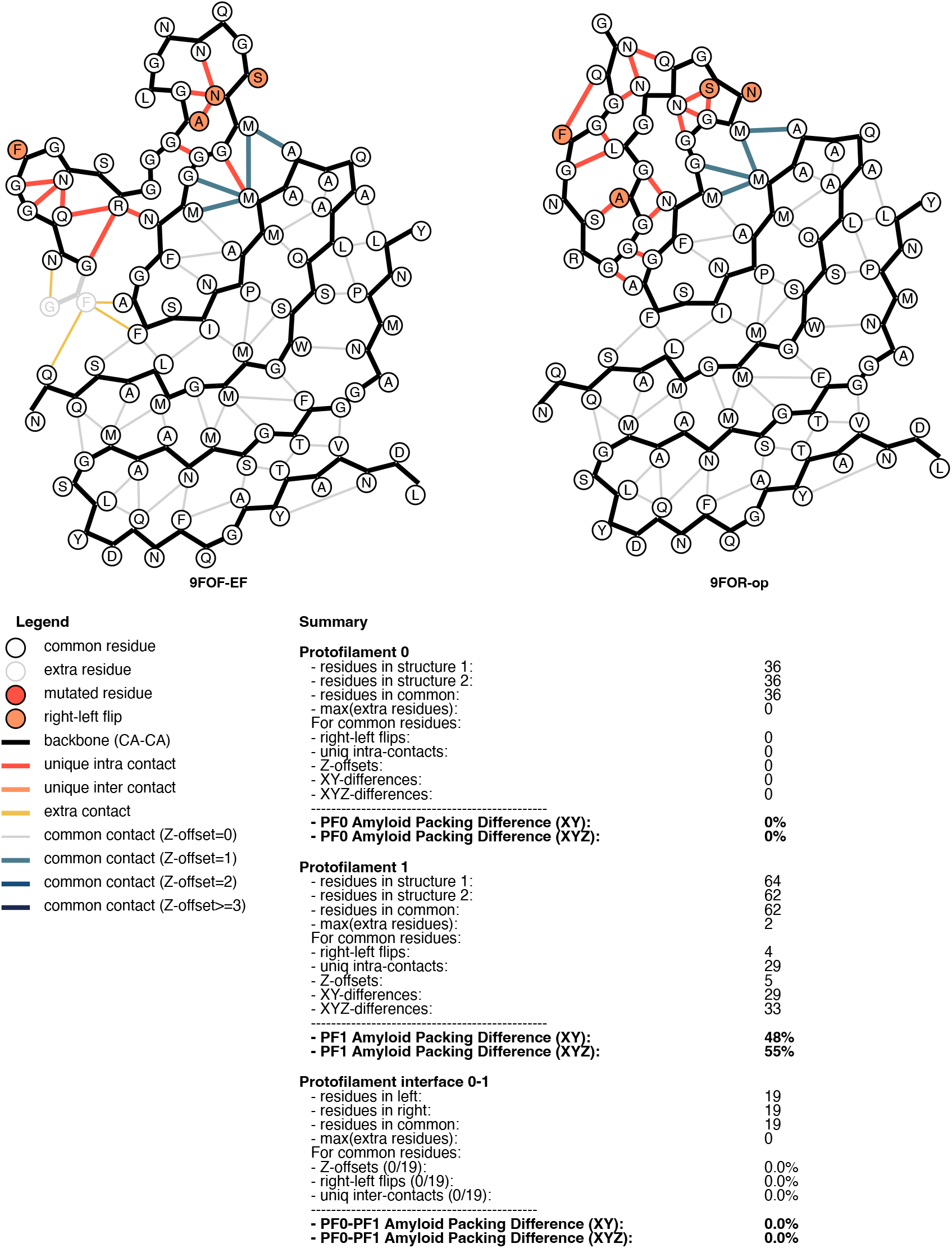
Output of the compare.py script for two variants of type C TDP-43+annexin A11 filaments. The ADP calculated over both protofilaments in PDB entries 9FOF and 9FOR is (29+2) / (64+36) = 31%.

